# Human REM sleep controls neural excitability in support of memory formation

**DOI:** 10.1101/2022.05.13.491801

**Authors:** Janna D. Lendner, Bryce A. Mander, Sigrid Schuh-Hofer, Hannah Schmidt, Robert T. Knight, Matthew P. Walker, Jack Lin, Randolph F. Helfrich

**Affiliations:** Hertie Institute for Clinical Brain Research, Center for Neurology, University Medical Center Tübingen; Hoppe-Seyler-Str 3, 72076 Tübingen, Germany; Department of Anesthesiology and Intensive Care Medicine, University Medical Center Tübingen; Hoppe-Seyler-Str 3, 72076 Tübingen, Germany; Department of Psychiatry and Human Behavior, UC Irvine; 101 The City Dr, Orange, CA 92868, USA; Department of Neurophysiology, University Medical Center Mannheim; Ludolf-Krehl-Str. 13-17, 68167 Mannheim, Germany; Department of Neurology and Epileptology, University Medical Center Tübingen; Hoppe-Seyler-Str 3, 72076 Tübingen, Germany; Helen Wills Neuroscience Institute, UC Berkeley; 130 Barker Hall, CA 94720, USA; Department of Psychology, UC Berkeley; 2121 Berkeley Way, CA 94720, USA; Department of Neurology, UC Davis; 3160 Folsom Blvd, Sacramento, CA 95816, USA; Center for Mind and Brain, UC Davis; 267 Cousteau Pl, Davis, CA 95618, USA

## Abstract

Sleep oscillations provide a key substrate to facilitate memory processing, the underlying mechanism of which may involve the overnight homeostatic regulation of plasticity at a synaptic and whole-network level. However, there remains a lack of human data demonstrating if and how sleep enhances memory consolidation and associated neural homeostasis. We combined intracranial recordings and scalp electroencephalography (EEG) in humans to reveal a new role for rapid eye movement (REM) sleep in promoting the homeostatic recalibration of optimal excitation/inhibition-balance. Moreover, the extent of this REM-sleep homeostatic recalibration predicted the success of overnight memory consolidation, expressly the modulation of hippocampal— neocortical excitability favoring remembering rather than forgetting. The findings describe a novel, fundamental role of human REM sleep in maintaining neural homeostasis, thereby enhancing long-term memory.

## Introduction

Contemporary theories of sleep function have proposed that the homeostatic regulation of excitability constitutes the physiologic mechanism of neural network plasticity facilitating overnight memory consolidation (Klinzing et al., 2019; Tononi and Cirelli, 2014). Numerous behavioral studies support a key role of sleep for stabilization of recently acquired memoires and restoring processing capacities for next day learning (Girardeau and Lopes-dos-Santos, 2021; Klinzing et al., 2019; Walker and Stickgold, 2006). Related, in vivo animals studies have revealed that wakefulness and learning lead to a progressive build-up of cellular synaptic potentiation and a net increase of excitability (Tononi and Cirelli, 2014; Vyazovskiy et al., 2008, 2009). In extreme, prolonged wakefulness using sleep deprivation in animal models impairs memory performance due to synaptic saturation (Huber et al., 2013; Weiss and Donlea, 2022). It has been proposed that, during sleep, a neural homeostatic process restores the optimal neural milieu for learning (Timofeev et al., 2001; Tononi and Cirelli, 2006, 2014). Specifically, accumulated daytime neural excitability is normalized through a process of enhanced neural inhibition and the elimination of synapses (termed ‘down-scaling’ or ‘pruning’). Through such mechanisms, it is proposed that the brain re-establishes the optimal excitatory/inhibitory (E/I) balance (Born and Feld, 2012; Grosmark et al., 2012; Vyazovskiy et al., 2009; Watson et al., 2016). That is, E/I-balance might constitute a marker of the degree of cellular and network plasticity during sleep (Tononi and Cirelli, 2006), though such a cellular-network proposal remains untested in humans, as does the possibility that this mechanism is functional and enhances human memory. The paucity of this knowledge relative to animal models exploring cellular synaptic pruning mechanisms (Li et al., 2017; Zhou et al., 2020) is a direct consequence of limited invasive recordings of neural processes in humans.

To date, the majority of the evidence for cellular and network homeostasis of neural excitability during sleep suggested that NREM slow oscillations (SO; < 1.25 Hz) constitute a key mechanism of homeostatic (down-) regulation (Tononi and Cirelli, 2006, 2014), since SOs periodically silence neural firing during ‘down-states’ (Steriade et al., 1993). Though less evidence exists, a similar role in rodent models has started to emerge for theta activity (4-10 Hz) during REM sleep (Grosmark et al., 2012). Unlike NREM SOs, which are common to both rodents and humans, only rodent REM sleep features prominent theta oscillations (Cantero et al., 2003), whereas human REM sleep is characterized by desynchronized electroencephalogram (EEG) activity without prominent oscillations. Therefore, traditional oscillations-centric approaches fail to capture homeostatic processes in human REM sleep; thus, there is a need to establish novel electrophysiological signatures of desynchronized, non-oscillatory brain activity.

In the electrophysiological power spectrum, oscillations are defined as distinct peaks that rise above the exponential decay function (1/f^*x*^ relationship between frequency and power; *x* reflects the spectral slope) of the background activity. Previously, background activity has often been discarded as neuronal noise (Voytek et al., 2015); however, recent findings revealed that it contains unique information about the underlying brain state (He, 2014; Lendner et al., 2020; Miller et al., 2009). Since background activity lacks a defining temporal scale, it has also been termed aperiodic or scale-free activity (Donoghue et al., 2020; He, 2014; Helfrich et al., 2021). Recently, novel computational models established that aperiodic brain activity captures the collective population activity of excitatory and inhibitory neurons (Chini et al., 2021; Gao et al., 2017). Specifically, increased inhibition results in a reduction (steepening of the EEG power spectrum) and excitation in an increase of aperiodic activity (flattening of the power spectrum; Chini et al., 2021; Gao et al., 2017). Furthermore, an important advantage of aperiodic features is that they can be obtained for every brain state including wakefulness from scalp and intracranial EEG; thus, providing a unique opportunity to link macro-scale signals to micro-scale properties (Chini et al., 2021; Gao et al., 2017; Kanth and Ray, 2020; Watson et al., 2018). Critically, recent findings revealed that REM sleep is associated with the most profound reduction of aperiodic activity during sleep (Lendner et al., 2020), i.e. reflecting an overall shift towards inhibition (Helfrich et al., 2021; Niethard et al., 2016). Thus, these novel assessment measures of aperiodic activity provide the opportunity to gain key insights into the function of human REM sleep, especially concerning mechanisms regulating network excitability and inhibition underlying neural homeostasis.

Aperiodic activity also tracks behaviorally relevant information, including arousal under general anesthesia, age-related cognitive decline or working memory performance (Donoghue et al., 2020; He, 2014; Lendner et al., 2020; Miller et al., 2009; Voytek et al., 2015). In addition, aperiodic activity has candidate properties to capture dynamics of overnight memory consolidation as its information-rich environment is optimal to imprint new mnemonic information onto existing circuits (Hanslmayr et al., 2016; Helfrich et al., 2021). While theoretical accounts suggested that homeostatic synaptic pruning could benefit memory consolidation (Boyce et al., 2017; Klinzing et al., 2019; Tononi and Cirelli, 2014), aperiodic activity now provides the necessary theoretical framework to link memory retention to the homeostatic regulation of neural activity in humans.

These considerations imply an important and to date unappreciated role of human REM sleep for neural homeostasis and memory retention. The current study tested five specific predictions. (1) Neural homeostasis of excitability during sleep should selectively counteract accumulated daytime excitation through increased inhibition. Thus, aperiodic activity as a marker of population E/I-balance should increase during the day (reflecting excessive excitation) and decrease after sleep, thereby, restoring the optimal E/I-balance for the next day. (2) Conversely, sleep loss will abate this homeostatic regulation, resulting in a surplus of excitation. (3) As REM sleep is characterized by increased inhibition concomitant with the strongest reduction of aperiodic activity (Lendner et al., 2020; Niethard et al., 2016), we hypothesized that REM sleep, rather then NREM sleep, mediates the overnight homeostatic control of excitability. If this downregulation is functionally relevant rather than epiphenomenal, then (4) the degree of aperiodic modulation should predict individual memory retention and (5) this modulation should preferentially occur in the neocortex, the key node for human long-term memory retention (Frankland and Bontempi, 2005).

We tested these predictions using invasive and non-invasive electrophysiological recordings in humans. We combined an overnight episodic memory task (Helfrich et al., 2018; Mander et al., 2013) with scalp EEG recordings of resting state before and after habitual sleep (study 1a; N = 40; **Fig. 1**), as well as after sleep deprivation (study 2; N = 12; **Fig. 1**) using novel tools for spectral parameterization to estimate population E/I-balance (Donoghue et al., 2020). Furthermore, we examined the aperiodic activity in overnight sleep recordings using either scalp EEG (study 1b; N = 40; **Fig. 2**) and simultaneous scalp and intracranial EEG recordings (study 3; N = 15; 498 bipolar contacts; **Fig. 3**) in patients with pharmacoresistant epilepsy that underwent invasive monitoring before surgical removal of the epileptic focus to access cortical and subcortical structures with high spatiotemporal resolution.

**Fig. 1.**
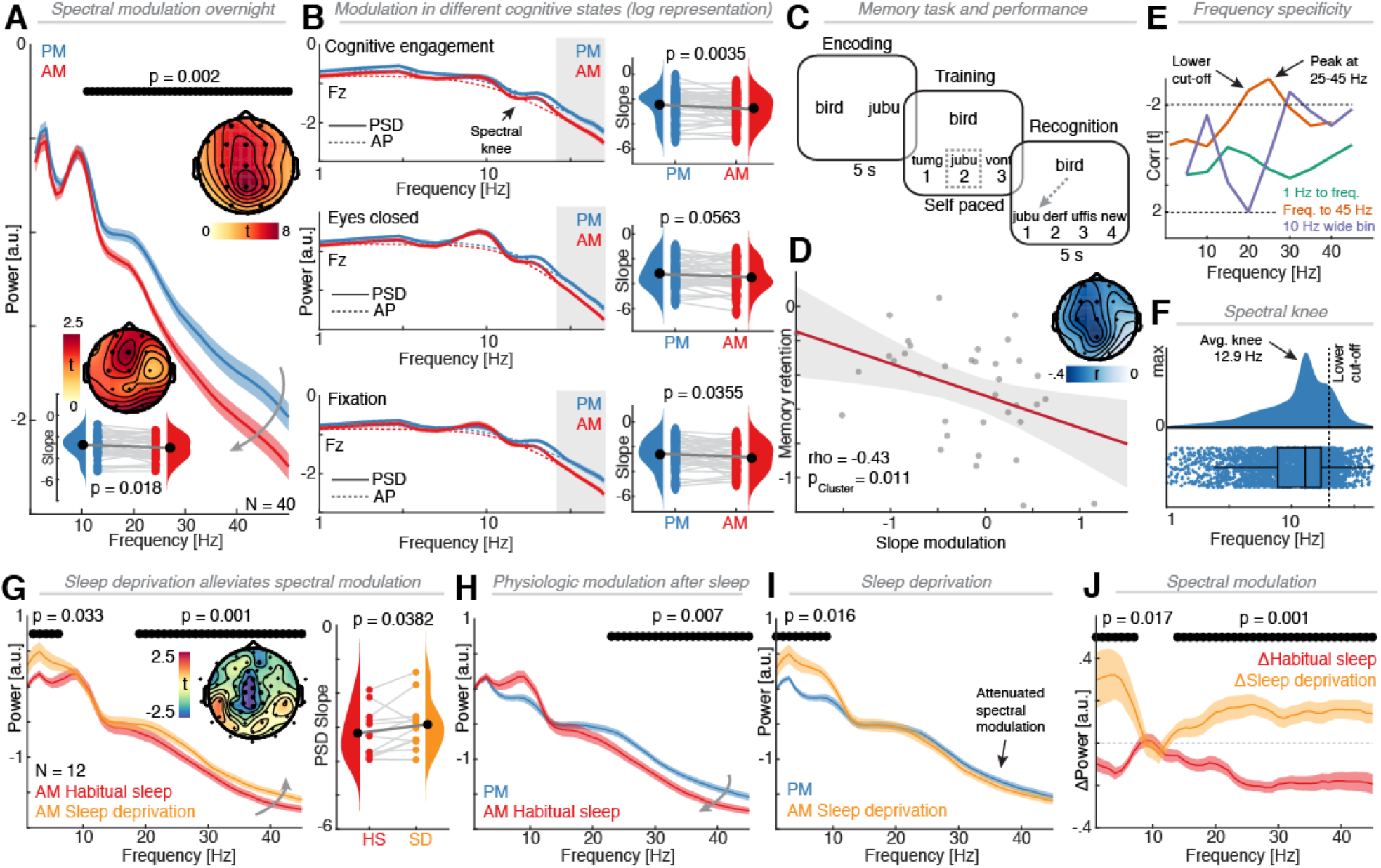
Overnight modulation of aperiodic activity predicts memory retention and is attenuated by sleep deprivation. (**A**) Grand average EEG power spectra before (PM) and after (AM) habitual sleep (N = 40; averaged across three conditions and all electrodes in cluster: cognitive engagement, eyes closed and central fixation; semi-log representation) indicate a broadband power decrease (11-50 Hz) over all electrodes (inset upper right) after sleep. Inset lower left: A decrease of spectral slope over frontal sensors (visualized at max. Fz) mirrors the broadband power decrement. (**B**) Left: Log-log spectral representation at Fz per cognitive state reveals that the power decrement is frequency-specific but state-invariant. Estimates of aperiodic background activity (dashed lines; FOOOF method) indicate a characteristic bend (‘knee’; black arrow) of the power spectrum. Right: State-invariant decreases of the spectral slope after sleep (Fz; paired one-tailed t-test). (**C**) Episodic word pair task. Participants learned 120 word-nonsense word associations. After encoding (left), participants were trained to criterion (center) prior to sleep and then performed recognition test prior and after sleep (right). (**D**) Cluster-corrected correlation analysis revealed a significant association between slope modulation (FOOOF model) and memory retention: Participants who showed a stronger slope decrease from PM to AM exhibited better memory retention. (**E**) The correlation analysis was repeated for different fit parameters (1st degree polynomial fitting; purple: 10 Hz wide fitting range with variable center frequencies (x-axis); green: fitting range with fixed start at 1 Hz and variable end point (x-axis); orange: fitting range with variable start (x-axis) and fixed end point at 45 Hz; visualized over midline EEG sensors) reveals the strongest EEG-behavioral correlation in the range from 25-45 Hz (inverted y-axis). Note that the correlation was reversed or non-significant if frequencies below the spectral knee were included (**Fig. S1F**). Correlation values were transformed to t-values, dashed lines indicate two-tailed p < 0.05 as derived from the inverse cumulative distribution function. (**F**) Distribution of spectral knees across all subjects, conditions and electrodes reveals a median knee frequency of 12.9 Hz (SD = 7.07 Hz; lower cut-off = 20 Hz (mean+1SD); cf. panel E). (**G**) Left: Eyes open resting state recordings in the AM reveal a broadband power increase after sleep deprivation (N = 12; cluster test: cluster 2-6 Hz, p = 0.04, d = 0.72; cluster 19-47 Hz, p = 0.0040; d = 0.87; visualized at Cz) over central sensors (inset), accompanied by an increase of spectral slope (right). (**H**) Comparison to pre-sleep eyes open recordings (PM) replicates the broadband down regulation by sleep from Study 1 (cf. panel A/B) while (**I**) sleep deprivation attenuates this effect. (**J**) Sleep-mediated physiologic broadband power decrease (red; AM vs. PM) and the adverse increase following sleep deprivation (orange; AM slept vs. AM sleep-deprived).

**Fig. 2.**
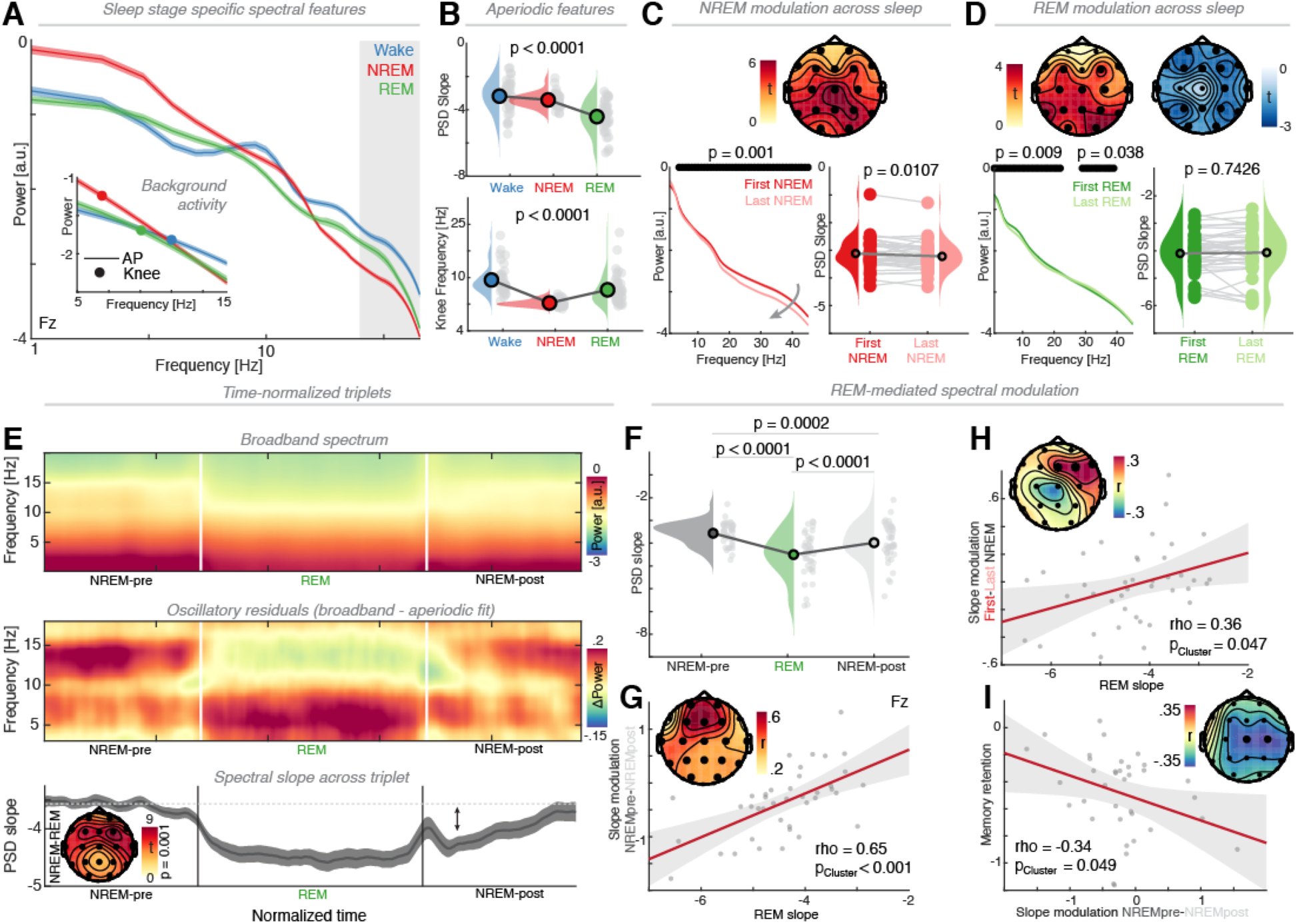
Aperiodic activity during REM sleep mediates overnight spectral modulation and memory retention. (**A**) Grand average power spectrum for different sleep stages. Grey shaded area indicates the peak fitting range (cf. **Fig. 1E**). Inset: Aperiodic activity and spectral knee. (**B**) Features of the aperiodic activity indicate a state-specific dissociation. (**C**) Spectral modulation across NREM sleep demonstrates a brain-wide (upper panel), broadband power reduction (lower left), which was well captured by a decrease of the spectral slope (lower right). (**D**) Spectral modulation across REM sleep shows a widespread (upper panels) frequency-specific (lower left) effect, which left the spectral slope unchanged. (**E**) Upper: Time-normalized triplet of adjacent NREM-REM-NREM segments (visualized at Cz). Center: Spectral residuals after subtraction of aperiodic activity reveal state-specific oscillatory signatures (NREM: spindle activity ∼14 Hz; REM: theta activity at ∼6 Hz; visualized in young adults to attenuate the spindle slowing-related spectral smearing). Lower: Spectral slope over time across all subjects. Note a decrement during REM sleep followed by a net decrease in subsequent NREM segment. Inset: Topographical depiction of slope differences between NREM and REM sleep reveal a frontal maximum (visualized at Fz). (**F**) Average spectral slopes across the triplet reveal significant time-dependent differences. (**G**) The spectral slope during REM sleep predicted the slope difference from NREM-pre to NREM-post (topographical inset depicts spatial extent), i.e. the steeper the slope during REM, the larger the down-modulation between the adjacent NREM segments. (**H**) A similar pattern was observed over frontal sensors, when the average REM slope was correlated against the difference between first and last NREM segment of the night (cf. panel C). (**I**) Confirming and extending the observation in **Fig. 1D**, a large down-regulation of slope across NREM sleep predicted better memory performance.

**Fig. 3.**
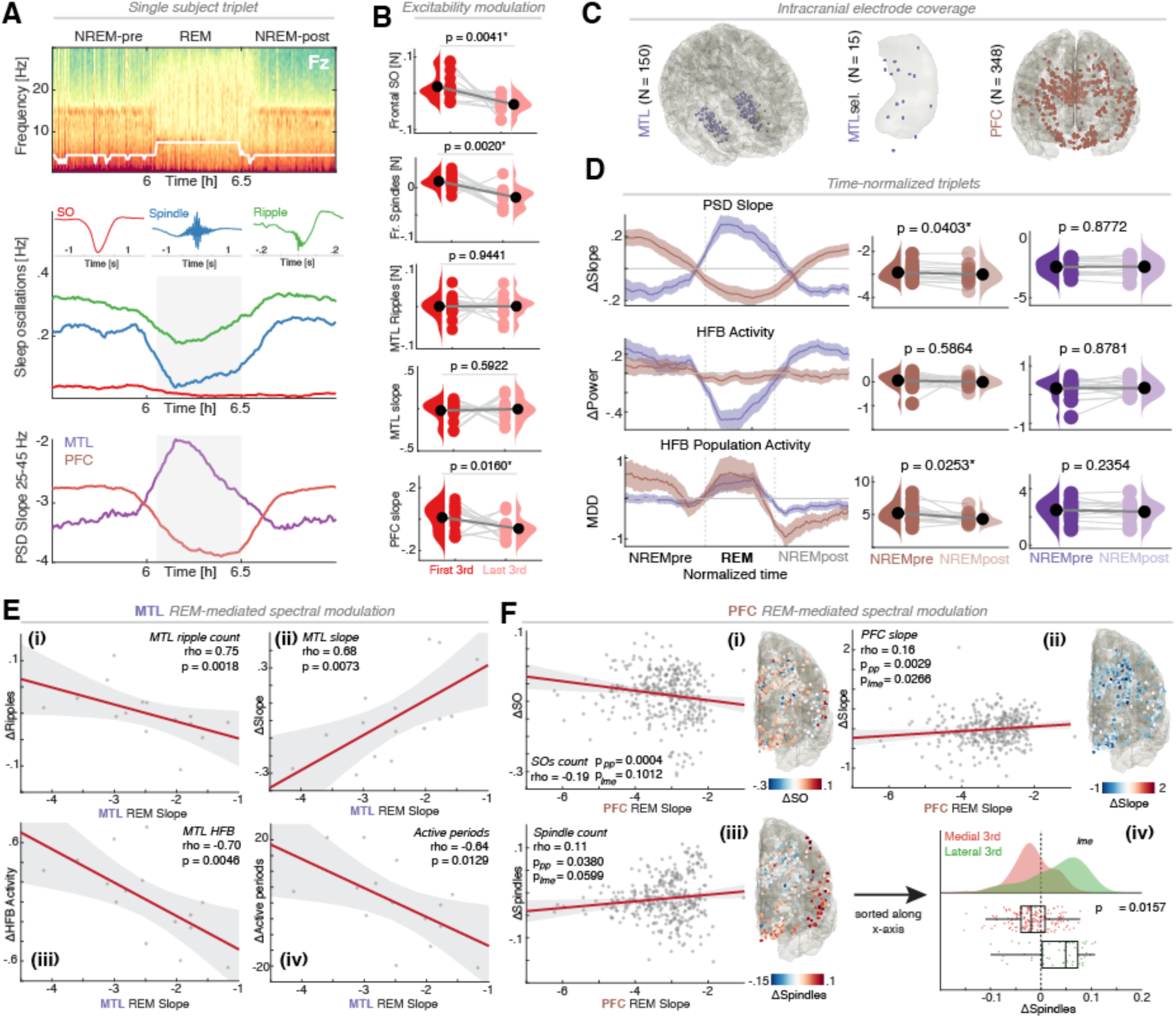
Aperiodic activity indexes sleep homeostasis and functionally dissociates medial temporal and prefrontal cortex. (**A**) Upper: Single subject example of NREM-REM-NREM triplet (visualized at Fz) of a participant with implanted electrodes in the MTL and PFC. Center: Mean waveform shapes of detected frontal SOs (red) and spindles (blue), as well as MTL ripples (green) and the rate modulation across the triplet with a prominent decrease during REM sleep. Lower: Average spectral slopes in the MTL and PFC across the triplet reveal a clear dissociation, with a flattening of the spectral slope in MTL during REM and a steepening in PFC. (**B**) Different surrogate markers indexing neuronal excitability across sleep highlight significant reductions in SO and spindle counts as well as frontal spectral slopes. (**C**) Electrode coverage across 15 participants with simultaneous coverage in the MTL and PFC. In every participant, we selected one MTL channel with the lowest number of epileptic discharges outside of the seizure onset zone for subsequent analyses. (**D**) Row 1: Spectral slope time-resolved across the triplet. Note the prominent, REM-mediated antagonistic modulation between PFC and MTL. We observed a reduction (steepening) of the spectral slope in NREM-post as compared to NREM-pre in PFC, but not MTL. Row 2: Average high frequency power (HFB; 120-200 Hz; above the ripple range) shows no modulation in PFC, and only little modulation across the triplet in MTL with no differences from pre-to post-NREM sleep. Row 3: Population activity (multidimensional distance (MDD) of the HFB trajectory in state space) highlights elevated population dynamics during REM sleep (i.e. dispersion of activity patterns), with a net decrease after REM sleep in the PFC. (**E**) MTL: Spearman rank correlations between REM slope difference (last 3rd – 1^st^ third, cf. panel B) and surrogate markers of neuronal excitability. A more negative slope during REM was associated with an increase in MTL ripple count (see i), high frequency power (see iii) and active periods (see iv), as well as a steepening of the PSD slope (ii). (**F**) The same analysis for the PFC showed a modulation of SO count (i), spindle count (iii) and PFC slope (ii) as a function of the REM slope at the given electrode (p values are reported for the Spearman correlation at the pseudo-population (pp) level as well as for a linear mixed effects model with subjects as random intercepts). Additionally, an inverse relationship between REM slope and spindle count was observed(iv), and was driven by prominent spatially specific differences. While the overall number of spindles was reduced over medial PFC, lateral and orbitofrontal PFC exhibited an increase across the night.

## Results

### Aperiodic activity is downregulated during sleep

To test whether homoeostatic downregulation of aperiodic activity occurred during sleep, the study employed resting state scalp EEG recordings in three cognitive states (N = 40; cognitive engagement during backward counting, eyes open central fixation, eyes closed quiescent rest) before and after a night of habitual sleep. Spectral analysis revealed a broadband power decrease after sleep in all frequencies above 11 Hz and across the majority of EEG sensors (**Fig. 1A**; averaged across all conditions, cluster test; p = 0.0020, d = 0.86). Importantly, this broadband modulation was solely driven by changes of non-oscillatory aperiodic brain activity (**Fig. S1A**).

Aperiodic activity was estimated from three parameters of the electrophysiological power spectrum: spectral slope *x* (the negative exponent of the 1/f^*x*^ decay function), y-intercept and the population time constant (the frequency where a bend/’knee’ occurs in the 1/f spectrum). Note that the slope and y-intercept provided redundant information (correlated at rho = -0.98), thus, subsequent analyses focused on the spectral slope. In line with a homeostatic downregulation of aperiodic activity, the spectral slope was more negative after habitual sleep, with the peak effect over frontal EEG sensors (inset **Fig. 1A**; cluster test; p = 0.0180, d = 0.32; Fz: PM: -2.76 ± 0.03, AM: -3.04 ± 0.03; mean ± SEM). Importantly, while there was a significant difference of the spectral slope between *cognitive states* (F_1.6,63.0_ = 33.32, p < 0.0001), all conditions exhibited a downregulation of aperiodic activity after sleep (**Fig. 1B**; 2-way RM-ANOVA at Fz; *PM/AM*: F_1,39_ = 6.35, p = 0.0159, *interaction*: F_2.0,77.8_ = 0.37, p = 0.69). To rule out potential confounds of muscle activity as a function of time of day, the analysis was repeated on the EMG. While muscle activity varied as a function of *cognitive state* (F_2.0,76.0_ = 5.07, p = 0.0086; *interaction*: F_1.7,65.2_ = 0.91, p = 0.3944), indicating more EMG-related activity in the two eyes open conditions, this was not the case for time of day (**Fig. S1B**; 2-way RM-ANOVA; *PM/AM*: F_1,38_ = 3.00, p = 0.0916; EMG was not available in one participant). Hence, subsequent analyses were performed on eyes closed data (highest signal-to-noise ratio across subjects; **Fig. S1C**). Taken together, these findings demonstrate that aperiodic activity undergoes a physiologic downregulation during the night.

### Homeostasis of aperiodic activity predicts successful memory retention

Having established a homeostatic modulation of aperiodic activity across sleep, we next investigated whether this modulation predicted long-term memory retention. Participants performed a sleep-dependent episodic memory test (**Fig. 1C**; 36 subjects completed behavioral testing). After encoding, participants were trained to criterion before initial recognition testing in the evening (Helfrich et al., 2018; Mander et al., 2013). After 8 h of sleep starting at their habitual bedtime, they performed the second recognition test the next morning. Participants who exhibited a stronger modulation of aperiodic activity (decrease of the spectral slope) demonstrated better memory retention (**Fig. 1D**; cluster test; p = 0.0011; mean rho = -0.36; **Fig. S1D**). This effect was not confounded by EMG activity or age (**Fig. S1E**; Spearman partial correlation at Fz: *EMG*: rho = -0.45, p = 0.0069; *age*: rho = -0.46, p = 0.0050). Next, we tested whether the correlation depended on the precise fitting range (first-degree polynomial fitting; **Fig. 1E**). Indeed, a significant correlation between behavior and slope was observed for center frequencies > 20 Hz (high congruence to initial parametrized estimates: Spearman rho = 0.99) with a peak in the range from 25-45 Hz. This effect was not biased by the presence of a spectral ‘knee’ below this frequency range (bend of the PSD at ∼12.9 Hz ± 7.1 Hz; median ± SD; reflecting a population time constant during wakefulness; **Fig. 1F**) where fits below or encompassing the knee frequency rendered the observed correlation non-significant (**Fig. 1E** and **Fig. S1F**). Collectively, this set of findings demonstrated that downregulation of aperiodic brain activity across the night is associated with memory retention.

### Sleep deprivation attenuates homeostatic regulation of aperiodic activity

Having characterized the spatiotemporal extent of the homeostatic regulation, we next assessed the causal role for sleep in the modulation of aperiodic activity across the night in an independent cohort that was sleep-deprived (N = 12; eyes open, central fixation). A disruption of habitual sleep resulted in a broadband power increase over central sensors (inset **Fig. 1G**; cluster test; p = 0.0030, d = 0.81), evident as a flattening of the spectral slope (post habitual sleep: -3.18 ± 0.20; sleep deprivation: -2.91 ± 0.22). Given the different electrode layouts between the two studies, all subsequent analyses were focused on the common central electrode Cz. First, the broadband power decrease after sleep was replicated (cf. **Fig. 1A**) using a between subject design (**Fig. 1H**; cluster test; p = 0.0070, d = 0.81). Statistical comparison to pre-sleep resting states revealed that sleep deprivation attenuated the observed modulation of aperiodic activity (**Fig. 1I**), and led to an increase of low frequency activity (cluster test; 1-9 Hz; p = 0.0200, d = 1.23, a finding in line with observations of slow waves after prolonged wakefulness (Vyazovskiy et al., 2011)). When directly contrasting the benefits of habitual sleep and adverse effects of sleep deprivation (**Fig. 1J**), a broadband shift between both conditions was observed (cluster test; cluster 14-48 Hz; p < 0.001, d = 1.50; cluster 1-7 Hz; p = 0.0180, d = 1.56). Taken together, these findings establish that causal manipulation of neural homeostasis by sleep deprivation attenuates the downregulation of aperiodic brain activity.

### Aperiodic activity in REM sleep recalibrates neural excitability

Next, we sought to determine if REM sleep, characterized by increased inhibition (Lendner et al., 2020) and implicated in reorganizing neuronal excitability (Grosmark et al., 2012), mediates downregulation of aperiodic activity. Theta oscillations have been implicated in maintaining network homeostasis in rodents, but are less prominent in humans than in rodents (**Fig. S2A/B**). Therefore, subsequent analyses focused on aperiodic activity, which had previously been shown to index inhibitory drive during REM sleep (Lendner et al., 2020). Specifically, we asked whether aperiodic brain activity during REM sleep can predict a homeostatic modulation from one NREM epoch to the next one as well as to overnight modulation (cf. **Fig. 1A**). Consistent with previous findings, the spectral slope was more negative during REM sleep compared to NREM and wakefulness (**Fig. 2A/B**; 1-way ANOVA: F_2.9,75.3_ = 61.78, p < 0.0001; wake: -3.18 ± 0.15; NREM: -3.41 ± 0.07; REM: -4.41 ± 0.15).

Importantly, a state-dependent modulation of the population timescale (Gao et al., 2020) as indexed by a change of knee frequency was observed (**Fig. 2B**; 1-way ANOVA: F_2.0,77.1_ = 32.21, p < 0.0001; wake: 9.88 ± 0.44 Hz; NREM: 6.45 ± 0.06 Hz;

REM: 8.42 ± 0.46 Hz). When contrasting the first and last NREM segment of the night, a broadband spectral power modulation was evident (**Fig. 2C**; p < 0.001, d = 0.80) with a similar spatial extent as the homeostatic effect across the night (cf. **Fig. 1A** and **Fig. S2C**). The broadband modulation had a steepening of the spectral slope (paired one-tailed t-test: t_39_ = 2.40; p = 0.0107, d = 0.38), but did not encompass the SO range (< 4 Hz). When directly contrasting the first and last REM episode of the night, however, modulations were band-limited (**Fig. 2D**; cluster test; cluster 1-23 Hz, p = 0.0090, d = 0.55; cluster 28-40 Hz, p = 0.0380, d = 0.39) and were not driven by a change in aperiodic activity (paired one-tailed t-test: t_39_ = -0.65, p = 0.7426, d = -0.10), resembling previous findings in rodents (Grosmark et al., 2012).

To determine if REM sleep mediates the modulation of aperiodic activity in subsequent NREM epochs, time-normalized triplets of NREM-REM-NREM sleep were extracted (**Fig. 2E**). State-specific oscillatory patterns (**Fig. 2E**, central panel) were only apparent after subtraction of aperiodic activity from composite power spectra (**Fig. 2E**, upper panel). Aperiodic activity, quantified as the spectral activity slope, was strongly modulated over the course of the triplet (**Fig. 2E**, lower panel). Statistical comparison of the spectral slope during REM and NREM sleep showed a brain-wide difference that peaked over frontal sensors (cluster test; p < 0.001, d = 0.97).

Consistent with a homeostatic influence of REM sleep on NREM, a greater negative spectral slope (i.e. a stronger downregulation of aperiodic activity) was observed in NREM epochs after a REM episode compared to before (**Fig. 2F**; paired two-tailed t-test; t_39_ = 4.04, p = 0.0002, d = 0.64). On an individual level, a more negative spectral slope in REM sleep predicted a stronger modulation across the triplet (**Fig. 2G**; cluster test; p = 0.001, mean rho = 0.44; peak correlation at Fz rho = 0.65). This relationship remained unchanged after accounting for theta power (partial correlation: rho = 0.59, p < 0.0001) and was also apparent when the REM slope was correlated against the individual difference between first and last NREM segments of the night (**Fig. 2H**; cluster test; p = 0.0470, mean rho = 0.36; cf. **Fig. 2C**). Moreover, this effect was not confounded by SO power (**Fig. S2D**; Spearman partial correlation; rho = 0.37, p = 0.0189) or REM theta power (rho = 0.34, p = 0.0349).

Importantly, this modulation reliably predicted individual memory performance (**Fig. 2I**; cluster test; p = 0.049, mean rho = -0.34), and became even more robust after accounting for theta power (partial correlation; rho = -0.49, p = 0.0025), thus, providing additional support for the behavioral relevance of changes in aperiodic activity (cf. **Fig. 1D**). Collectively, these observations indicate that aperiodic activity tracks population E/I-balance across different brain states. Specifically, aperiodic activity during REM sleep predicts sleep-dependent homeostasis across the night with stronger reduction of aperiodic activity leading to superior memory retention.

### Distinct aperiodic activity regimes govern REM sleep in MTL and PFC

Regional population E/I-balance was assessed in intracranial recordings (N = 15) in two key nodes of the human memory network, the prefrontal cortex (PFC) and the medial temporal lobe (MTL). Contemporary theoretical frameworks posit that long-term memory consolidation is associated with human PFC plasticity (Frankland and Bontempi, 2005; Klinzing et al., 2019) and we tested if the homeostatic modulation differentially impacts PFC and MTL. Analogous to previous work (Grosmark et al., 2012; Watson et al., 2016), a set of parameters was obtained, including the number of frontal SOs, frontal spindles and hippocampal ripples as well as spectral slopes, high frequency power and active episodes (Methods) in the MTL and PFC (**Fig. 3A**).

Across the night (comparison of NREM thirds; **Fig. 3B**), the count of SOs (1-way RM-ANOVA across all thirds; F_1.6,22.2_ = 8.01, p = 0.0041, d = 0.90), spindles (F_1.7,24.4_ = 8.76, p = 0.0020, d = 1.10) and the PFC slope (F_1.8, 25.6_, p = 0.0160, d = 0.57; all other markers p > 0.21) showed significant decreases. Subsequently, triplets for all subjects were extracted separately for MTL and PFC (**Fig. 3C**). A state-and region-specific modulation of the spectral slope was observed (**Fig. 3D**) with a prominent functional dissociation between the MTL and PFC (**Fig. 3D**; 2-way RM-ANOVA; ROI: F_1,14_ = 9.38, p = 0.0084; stage: F_1.2,17.1_ = 0.64, P = 0.4672; interaction: F_1.2,17.7_ = 21.95, p = 0.0001).

This analysis revealed a steepening of the PFC power spectrum during REM sleep (cf. **Fig. 2E**; paired two-tailed t-test; t_14_ = 3.44, p = 0.0039, d = 0.89) as well as a net decrease in aperiodic activity after a REM episode (replication of **Fig. 2F**; t_14_ = 2.26, p = 0.0403, d = 0.58). Critically, the pattern was reversed in the MTL (increase of the spectral slope during REM; t_14_ = -4.68, p = 0.0004, d = 1.21) and no REM-mediated modulation was observed (t_14_ = -0.16, p = 0.8772, d = 0.04).

In sum, these results reveal a double dissociation between the PFC and MTL REM sleep with increased inhibition observed during neocortical PFC indexed REM sleep providing the optimal neurophysiological milieu to induce neuroplasticity in support of long-term memory retention. In contrast to neocortical inhibition, a switch to more excitable dynamics was evident in the MTL, where no REM-mediated homeostatic down-regulation was observed.

Analysis of high frequency band (HFB) activity, a marker of multi-unit firing and dendritic synaptic potentials (Leszczyński et al., 2020), confirmed the observation of enhance hippocampal excitability. Mean HFB activity only changed in MTL but not PFC over the course of the triplet (*MTL*: F_1.3,18.9_ = 15.94, p = 0.0003; *PFC*: F_1.5,21.7_ = 0.39, p = 0.6279) with no homeostatic modulation (both p-values > 0.59). However, we found a dispersion of activity patterns across all recording sites (**Fig. 3D**, last row). Population vector analysis (Ebitz and Hayden, 2021) revealed a regionally-specific modulation of the multidimensional distance across the triplet (2-way RM-ANOVA; *ROI*: F_1,14_ = 42.07, p < 0.0001, *state*: F_1.2,16.6_ = 3.77, p = 0.0634; *interaction*: F_1.5,21.3_ = 1.84, p = 0.1878), which reflects a more heterogeneous and less synchronized population response. Importantly, we again observed homeostatic downscaling following REM sleep in PFC (t_14_ = 2.50, p = 0.0253, d = 0.65) but not in the MTL (t_14_ = 1.24, p = 0.2354, d = 0.32). To further quantify the REM-mediated modulation, we separately correlated the overall REM slope (analogous to **Fig. 2G/H**) with surrogate excitability markers.

Larger negative slopes during REM sleep predicted increased overnight hippocampal ripple activity (Spearman rho = -0.75, p = 0.0018), HFB activity (rho = - 0.70, p = 0.0046) and active periods (rho = -0.64, p = 0.0129). This was observable on the individual subject level (**Fig. 3E**), including a steepening of the spectral slope across the night (**Fig. 3E**; rho = 0.68, p = 0.0073; replicating **Fig. 2H**). Importantly, the same relationship between REM and the overnight slope decrease was observed in PFC (**Fig. 3F**; rho = 0.16, p_pseudo-population_ = 0.0029, p_lme_ = 0.0266; t_345_ = 2.23; CI_95_ = [0.007 0.109]; p-values were calculated in a pseudo-population and confirmed using a linear mixed effects models with subjects as random intercepts). In addition, the count of prefrontal SO (rho = -0.19, p_pseudo-population_ = 0.0004, p_lme_ = 0.1012; t_345_ = 1.64; CI_95_ = [-0.001 0.012]) and spindles changed as a function of REM slope (rho = 0.11, p_pseudo-population_ = 0.0380, p_lme_ = 0.0599; t_345_ = -1.89; CI_95_ = [-0.009 0.0002]). The spindle modulation exhibited an opposite pattern between medial and lateral frontal cortex, with a decrease in medial and an increase in lateral prefrontal regions (Fig. 3F; p_lme_ = 0.0157; t_345_ = 2.43; CI_95_ = [5×10^−5^ 5×10^−4^]).

Together, these results reveal that aperiodic activity during REM sleep predicts homeostatic modulation of neural excitability in sleep. There was a region-specific modulatory influence on NREM oscillations: while homeostatic modulation of aperiodic activity in the MTL predicted a modulation of hippocampal ripples, neocortical activity predicted a modulation of SOs and spindles. Critically, the post-REM excitability modulation was confined to the neocortex indicating that REM-mediated homeostatic downregulation preferentially impacts neocortical regions to support long-term memory retention.

## Discussion

Our results from three independent studies demonstrated that REM sleep mediates sleep-dependent neural homeostasis by downregulation of excitability in humans, which predicted the success of subsequent overnight long-term memory retention. While previous work has primarily focused on sleep oscillations (Klinzing et al., 2019; Miyawaki and Diba, 2016; Tononi and Cirelli, 2014), and largely those of NREM sleep, these results reveal that non-oscillatory, aperiodic activity during REM sleep is an inherent characteristic of the functional organization of the sleeping human brain. Unlike state-specific sleep oscillations, aperiodic activity can be estimated for every state including wakefulness constituting a marker to directly compare activity across different neural and behavioral states. A causal manipulation of neural homeostasis through the removal of sleep using total overnight deprivation resulted in an increase in excitation and an attenuated downregulation of aperiodic activity. Furthermore, REM sleep mediated overnight neural homeostasis by down-regulating aperiodic activity to establish optimal excitation/inhibition-balance. In addition, it was during REM sleep that a pronounced functional and anatomical dissociation was observed between two key brain regions for memory, the MTL and neocortex. Specifically, the MTL switched from a stable inactive state in NREM sleep to a transient active state during REM sleep, while the neocortex transitioned from an active to an inactive state. Moreover, REM sleep mediated the homeostatic regulation of oscillatory NREM sleep signatures in a spatially specific manner, with aperiodic activity in the MTL indexing the modulation of hippocampal ripples, while neocortical aperiodic activity predicted SO and spindle modulation. These findings indicate a novel interactions between sleep-stages, such that the expression of NREM sleep oscillations are governed as a function of the modulation of population level excitatory versus inhibitory balance by the preceding REM sleep episode.

### E/I-balance tracks neural homeostasis during sleep

Returning to the fundamental experimental hypothesis, exactly how does the sleeping brain regulate neural homeostasis to meet the demands of optimal function, including that required for information processing and memory retention? During wakefulness and learning, new synapses are formed, existing connections are strengthened and overall neural firing increases, thus, resulting in an accumulation of excitation with time spent awake (Maret et al., 2011; Vyazovskiy et al., 2008, 2009). Sleep has been proposed to counteract this progressive build-up of excitation to maintain healthy neural functioning, with sleep deprivation attenuating such homeostatic regulation and impairing cognitive processes and memory formation (Huber et al., 2013; Vyazovskiy et al., 2009; Weiss and Donlea, 2022).

On the cellular level, sleep reduces neural firing (Vyazovskiy et al., 2009) and promotes synapse elimination (Maret et al., 2011; Vyazovskiy et al., 2008) to rebalance excessive daytime excitation. At the population level, strong inhibitory bouts occur as ‘down states’ during sleep, which manifest as SOs in meso- and macro-scale recordings (Steriade et al., 1993; Timofeev et al., 2001). Hence, one seminal hypothesis posits that SO-mediated post-synaptic depression counteracts excitation to restore the optimal E/I-balance (Tononi and Cirelli, 2006, 2014). While SOs and related NREM sleep oscillations exhibit several candidate properties (Miyawaki and Diba, 2016; Norimoto et al., 2018), how the homeostatic excitability regulation as observed on the cellular level translates to macro-scale recordings at the in vivo level in the human cortex it poorly understood, despite being fundamental for affirming sleep-dependent neural network recalibration and its functional consequences.

Novel computational models have proposed a missing link between cellular and macro-scale signals (Chini et al., 2021; Gao et al., 2017). Specifically, aperiodic activity as indexed by the spectral exponent of the PSD predicts the E/I-balance of the underlying neuronal population (Gao et al., 2017; Helfrich et al., 2021; Lendner et al., 2020). The present findings tested the prediction that aperiodic activity is homeostatically regulated during sleep. While physiologic sleep increased inhibition (decreased the spectral exponent), sleep deprivation attenuated the homeostatic regulation (increased the spectral exponent). In contrast to the popular notion that inhibition is maximized during NREM sleep given the ubiquitous expression of SOs (Steriade, 2003), the strongest inhibition was observed during REM sleep. This observation raises the question if REM sleep inhibition mediates the overnight recalibration of population E/I-balance in humans.

### REM sleep inhibition recalibrates the E/I-balance during sleep

While SOs during NREM sleep have typically been linked to inhibition (Steriade, 2003), mounting evidence suggests that such NREM sleep consequences are nuanced, and also reflect a brain state of considerable excitability (Klinzing et al., 2019; Tamaki et al., 2020; Watson et al., 2016). For example, NREM sleep increases synaptic strength and neural firing at a cellular level (Yang et al., 2014). At the population level, the cardinal oscillations of NREM sleep actively coordinate the hippocampal-neocortical dialogue to enable information reactivation, transfer and consolidation (Buzsáki, 2015; Klinzing et al., 2019). Thus, NREM sleep oscillations, including SOs, have been suggested to mediate neuroplasticity through repetitive replay of firing sequences (Buzsáki, 2015; Ólafsdóttir et al., 2018) and the memory-specific upregulation of synapse formation (Yang et al., 2014), thus reflecting a potential state of *increased* net excitation, in addition to co-occurring benefits of synaptic downscaling (Tononi and Cirelli, 2006, 2014).

In contrast, emerging evidence in animal models indicates a possible neuronal *inhibitory* state for REM sleep (Timofeev et al., 2001). At a cellular level, REM sleep promotes global synapse elimination (Li et al., 2017; Zhou et al., 2020). Moreover, two-photon calcium imaging (Niethard et al., 2016) and in-vivo electrophysiology studies (Grosmark et al., 2012; Watson et al., 2016) report a global reduction of neural firing with a selective increase of inhibitory cell activity during REM sleep, in accord with macro-scale findings demonstrating a reduction of aperiodic activity reflecting net inhibition (Lendner et al., 2020). The present results provide direct in vivo evidence corroborating this proposal in human cortex, showing that REM-mediated inhibition recalibrates excitability dynamics of the brain during this sleep state (Grosmark et al., 2012; Helfrich et al., 2021; Placidi et al., 2013; Watson et al., 2016). First, aperiodic REM activity predicted the homeostatic downregulation of the E/I-balance across the night. Second, REM sleep mediated a spatially specific homeostatic regulation of NREM sleep oscillations, such as SOs, spindles and ripples. Collectively, these findings reveal that homeostatic control of excitability is a core function of human REM sleep.

### REM sleep-mediated neural homeostasis predicts memory retention

Is the change in E/I-balance during REM sleep epiphenomenal, or instead, functional, specifically regarding sleep-dependent overnight memory processing (Born and Feld, 2012)? At the cellular level, consolidation of mnemonic representations requires a selective, activity-dependent elimination of synapses (Tononi and Cirelli, 2006). As this ‘down-scaling’ occurs primarily in sleep, prolonged wakefulness is proposed to result in hyper-excitability and synapse saturation leading to impaired memory function (Basner et al., 2013; Bridi et al., 2020; Vyazovskiy et al., 2009). Consistent with this proposition, when inhibitory cells are optogenetically inactivated during REM sleep in rodents, neural excitability increases and memory formation is impaired (Boyce et al., 2016). Conversely, REM sleep deprivation in rodents reduced synaptic plasticity (McDermott et al., 2006).

This set of findings suggests a role for REM sleep in maintaining neural E/I-balance in support of memory retention. The present scalp and intracranial cortical recordings in humans confirm that REM sleep mediates the downregulation of cortical excitability during the night. This recalibration benefit from reduced neural excitability had a functional benefit predicting successful next day memory retention. Indeed, this memory enhancement association was regionally specific to a local cortical network, that of the prefrontal cortex, in line with the idea that neocortical areas house long-term mnemonic storage (Frankland and Bontempi, 2005).

## Conclusions

Collectively, these findings uncover a fundamental role of human REM sleep in maintaining the neural homeostasis between excitation and inhibition, one that supports long-term memory formation. Our results establish that the homeostatic recalibration association with REM sleep activity constitutes an inherent and previously unidentified functional property of the sleeping brain, which dynamically controls experience-dependent excitability to enhance memory consolidation.

## Funding

German Research Foundation, Emmy Noether Program, DFG HE8329/2-1 (RFH) German Research Foundation, DFG LE 3863/2-1 (JDL)

Hertie Foundation, Network for Excellence in Clinical Neuroscience (RFH)

Jung Foundation for Research and Science, Ernst Jung Career Advancement Award in Medicine (RFH)

German Research Foundation, SFB1158 Association Project (SSH, RFH)

National Institutes of Health (R01AG031164, RF1AG054019, RF1AG054106 (MPW) National Institute of Health 2 RO1 NS021135 (RTK)

National Institutes of Health U19NS107609 (JJL, RTK)

## Author contributions

Conceptualization: JDL, RFH Methodology: JDL, RFH

Investigation: JDL, BAM, SSH, HS, RTK, MPW, JJL, RFH

Visualization: JDL, RFH

Funding acquisition: RTK, MPW, RFH Project administration: RFH Supervision: RFH

Writing – original draft: JDL, RFH

Writing – review & editing: BAM, SSH, RTK, MPW, JJL

## Data and materials availability

Freely available software and algorithms used for analysis are listed where applicable. All custom scripts and data contained in this manuscript are available upon request from the Lead Contact. Data will be deposited on Dryad upon acceptance.

## Materials and Methods

### Participants

Study 1: 26 healthy older (73.0 ± 5.4 years; mean ± SD) and 14 younger adults (20.6 ± 2.2 years; mean ± SD) participated in the study. All participants provided written informed consent according to the local ethics committee (Berkeley Committee for Protection of Human Subjects Protocol Number 2010-01-595) and the 6th Declaration of Helsinki. Here we report a subset of participants from a larger cohort that completed three resting state recordings in addition to overnight sleep recordings(Helfrich et al., 2018; Mander et al., 2013).

Study 2: 12 young healthy controls (mean age: 23.2 ± 1.1 years; seven men, five women) participated in the study. All participants provided written informed consent according to the local ethics committee at the University of Mannheim (Protocol number 2010-311N-MA) and the 6th Declaration of Helsinki. The resting state data was acquired in the context of a larger study investigating the effects of sleep deprivation on habituation, but have not been reported previously (Schuh-Hofer et al., 2015).

Study 3: We obtained intracranial recordings from 15 pharmacoresistant epilepsy patients (35.0 ± 11.1 years; mean ± SD; 9 female) who underwent pre-surgical monitoring with implanted depth electrodes (Ad-Tech), which were placed stereo-tactically to localize the seizure onset zone. All patients were recruited from the University of California Irvine Medical Center, USA. Electrode placement was exclusively dictated by clinical considerations and all patients provided written informed consent to participate in the study. Patients selection was based on MRI confirmed electrode placement in the MTL and PFC from a larger cohort (Helfrich et al., 2019; Lendner et al., 2020). We only included patients where one seizure free night was available and a sufficient amount of REM sleep was recorded (see inclusion criteria below). The study was not pre-registered. All procedures were approved by the Institutional Review Board at the University of California, Irvine (protocol number: 2014-1522) and conducted in accordance with the 6th Declaration of Helsinki.

### Experimental design and procedure

Study 1: All participants were trained on the episodic word-pair task in the evening and performed a short recognition test after 10 min. Then, participants were offered an 8 h sleep opportunity, starting at their habitual bedtime (**Table S1**). Resting state recordings were obtained directly before and after sleep. Polysomnography was collected continuously. Participants performed a long version of the recognition test approximately 2 h after awakening. Subsequently, we obtained structural MRI scans from all participants. Two older adults did not complete behavioral testing, and two young adults failed to achieve criterion at encoding. Thus, these four subjects were excluded from behavioral analyses, but were included in all electrophysiological analyses.

Study 2: In the three days prior to the experiment, sleep was monitored using an Actiwatch Device (Philips Respironics, Amsterdam). Participants were randomly assigned to either start in the sleep deprivation or habitual sleep group. In the experimental night, participants were either allowed to sleep and monitored using the Actiwatch device or kept awake and engaged by an experimenter. Recordings were obtained in the late AM or around noon.

Study 3: We recorded a full night of sleep for every participant. Recordings typically started around 8.00-10.00pm and lasted for ∼10-12h (**Table S2**). Only nights that were seizure-free were included in the analysis. Polysomnography was collected continuously.

### Behavioral task

Study 1: We utilized a previously established sleep-dependent episodic memory task (Figure 1A), where subjects had to learn word-nonsense word pairs (Mander et al., 2013). In brief, words were 3-8 letters in length and drawn from a normative set of English words, while nonsense words were 6-14 letters in length and derived from groups of common phonemes. During encoding, subjects learned 120 word-nonsense pairs. Each pair was presented for 5 s. Participants performed the criterion training immediately after encoding. The word was presented along with the previously learned nonsense word and two new nonsense words. Subjects had to choose the correctly associated nonsense words and received feedback afterwards. Incorrect trials were repeated after a variable interval, and were presented with two additional new nonsense words to avoid repetition of incorrect nonsense words. Criterion training continued until correct responses were observed for all trials.

During recognition, a probe word or a new (foil) probe word was presented along with 4 options: (1) the originally paired nonsense word, (2) a previously displayed nonsense word, which was linked to a different probe (lure), (3) a new nonsense word or (4) an option to indicate that the probe is new. During the recognition test after a short delay (10 min), 30 probe and 15 foil trials were presented. At the long delay (10 h), 90 probe and 45 foil trials were tested. All probe words were presented only once during recognition testing, either during short or long delay testing.

### Sleep monitoring and EEG data acquisition

Study 1: Polysomnography (PSG) sleep monitoring was recorded on a Grass Technologies Comet XL system (Astro-Med), including 19-channel electroencephalography (EEG) placed using the standard 10-20 system as well as Electromyography (EMG). Electrooculogram (EOG) was recorded the right and left outer canthi. EEG recordings were referenced to bilateral linked mastoids and digitized at 400 Hz. Sleep scoring was performed according to standard criteria in 30 s epochs (Rechtschaffen and Kales, 1968). Non-REM sleep (NREM) was defined as NREM stages 2-4. First and last NREM and REM epochs were defined as the first and last five minutes of the respective stages in the hypnogram.

Study 2: Resting state EEG recordings were obtained using a 64-channel BrainAmp amplifier (Brain Products GmbH) EEG system with equidistant Ag–AgCl electrode positions (EasyCap, Herrsching, Germany). The central electrode of this layout corresponded to electrode Cz (10-20 layout) and was therefore used for between group comparisons.

Study 3: We recorded from all available intracranial electrodes. In order to facilitate sleep staging based on established criteria, we also recorded scalp EEG, which typically included recordings from electrodes Fz, Cz, C3, C4 and Oz according to the international 10-20 system. Electrooculogram (EOG) was recorded from 4 electrodes, which were placed around the right and left outer canthi. All electrophysiological data was acquired using a 256-channel Nihon Kohden recording system (model JE120A), analog filtered at 0.01 Hz and digitally sampled at 5000 Hz. All available artifact-free scalp electrodes were low-pass filtered at 50 Hz, demeaned and de-trended, down-sampled to 400 Hz and referenced against the average of all clean scalp electrodes. EOGs were typically bipolar referenced to obtain one signal per eye. A surrogate electromyogram (EMG) signal was derived from electrodes in immediate proximity to neck or skeletal muscles, by high-pass filtering either the ECG or EEG channels above 40 Hz. Sleep staging was carried out according to Rechtschaffen and Kales guidelines by trained personnel in 30 second segments (Rechtschaffen and Kales, 1968) as reported previously (Helfrich et al., 2018; Mander et al., 2013). Same conventions as in study 1 were used.

### CT and MRI data acquisition

Study 3: We obtained anonymized postoperative CT scans and pre-surgical MRI scans, which were routinely acquired during clinical care. MRI scans were typically 1mm isotropic.

## Quantification and statistical analysis

### Behavioral data analysis

Study 1: Memory recognition was calculated by subtracting both the false alarm rate (proportion of foil words, which subjects’ reported as previously encountered) and the lure rate (proportion of words that were paired with a familiar, but incorrect nonsense word) from the hit rate (correctly paired word-nonsense word pairs). Memory retention was subsequently calculated as the difference between recognition at long minus short delays.

### EEG data

#### Preprocessing

Study 1/2 - Resting state: EEG data were imported into MATLAB and analyzed using the FieldTrip toolbox. Raw recordings were demeaned, detrended, high-pass filtered at 1 Hz, common average referenced and epoched into three-second-long segments with 50% overlap. Artifact detection was done semi-automatically for EOG, jump and muscle artifacts and visually confirmed (Oostenveld et al., 2011).

Study 1 - Sleep: EEG data were imported into FieldTrip, then demeaned, detrended, common average referenced and epoched into non-overlapping 30 second segments. Artifact detection was done manually in five-second segments (Helfrich et al., 2018).

Study 3: Scalp EEG was demeaned, de-trended and locally referenced against the mean of all available artifact-free scalp electrodes. We applied a 50 Hz low-pass filter and down-sampled the data to 500 Hz. All scalp EEG analyses were done on electrode Fz. In a subset of subjects Fz was not available and Cz was utilized instead of Fz. Intracranial EEG: In every subject, we selected all available electrodes in the medial temporal lobe, which were then demeaned, de-trended, notch-filtered at 60 Hz and its harmonics, bipolar referenced to its immediate lateral neighboring electrode and finally down-sampled to 500 Hz. We retained all MTL channels, but discarded noisy PFC channels. We adopted a previously introduced approach where we first detected interictal epileptic discharges using automated detectors (see below), which were then excluded from further analysis. Finally, we selected one MTL electrode per participant with the lowest number of overall detections. For PFC analyses, all available contacts in these regions were included and the same pre-processing steps were applied. Then all resulting traces were manually inspected and noisy, epileptic and artifact-contaminated PFC channels were excluded.

#### Extraction of REM epochs and time normalization procedure

REM epochs were detected based on the manually staged hypnogram (Grosmark et al., 2012). We first detected all REM epochs and then selected artifact-free epochs that spanned at least three consecutive epochs (90s) and required that the majority of adjacent periods within a time window ± 9 min were staged as NREM sleep. Subsequently, the identified REM epochs were extracted as continuous time-domain signals, then epoched into 100 overlapping epochs and subjected to multi-taper spectral analysis as outlined below. Similarly, we the adjacent NREM data were epoched into 10 second long segments with 70% overlap. The spectral estimates were then concatenated to form the final time-normalized triplet in the frequency domain. For statistical testing, we omitted the transition states and selected one third of the time-normalized epoch (beginning, center and end of the triplet, respectively) for subsequent testing.

#### Spectral analysis

Scalp EEG: Resting state spectral estimates were obtained through multitaper spectral analyses (Mitra and Pesaran, 1999; Prerau et al., 2017), based on discrete prolate slepian sequences. Spectral estimates were obtained between 1 and 50 Hz in 1 Hz steps. We adapted the number of tapers to obtain a frequency smoothing of ± 2 Hz.

Intracranial EEG: Spectral estimates were by means of multitaper spectral analyses based on discrete prolate spheroidal sequences in 153 logarithmically spaced bins between 0.25 and 181 Hz (Mitra and Pesaran, 1999). We adjusted the temporal and spectral smoothing to approximately match a ± 2 Hz frequency smoothing.

#### Estimation of aperiodic background activity

FOOOF fitting: In order to obtain estimates of aperiodic background activity, we first used the FOOOF algorithm (Donoghue et al., 2020). EEG spectra were fitted in the range from 1 to 45 Hz. Aperiodic background activity was defined by its slope parameter χ, the y-intercept *c* and a constant *k* (reflecting the knee parameter).

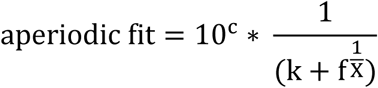

The relationship of the knee parameter and the knee frequency is given by

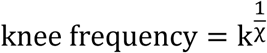

If a knee parameter could not be determined, we re-fitted the spectrum in the fixed mode, which is equivalent to a linear fit where k = 0.

Polynomial fitting: To estimate the spectral slope in different frequency bands, we also utilized first-degree polynomial fitting (Lendner et al., 2020), thus, yielding an instantaneous spectral exponent (slope, χ) and offset (y-axis intercept, *c*), for a given fitting range. EEG spectra were fitted using variable endpoints (from 1 Hz to 5-45 Hz, 5 Hz steps), variable starting points (to 45 Hz, from 5-40 Hz, 5 Hz steps) or a fixed bandwidth with varying center frequencies (5-45 Hz; ± 5 Hz).

#### Event detection

Slow oscillations (SO): Event detection was performed for every channel separately based on previously established algorithms (Helfrich et al., 2018; Staresina et al., 2015). We first filtered the continuous signal between 0.16 and 1.25 Hz and detected all the zero crossings. Then events were selected based on time (0.8 – 2 s duration) and amplitude (75% percentile) criteria. Finally, we extracted 5 s long segments (± 2.5 s centered on the trough) from the raw signal and discarded all events that occurred during an IED.

Sleep spindles: Based on established algorithms (Helfrich et al., 2018; Staresina et al., 2015), we filtered the signal between 12-16 Hz and extracted the analytical amplitude after applying a Hilbert transform. We smoothed the amplitude with a 200 ms moving average. Then the amplitude was thresholded at the 75% percentile (amplitude criterion) and only events that exceeded the threshold for 0.5 to 3 s (time criterion) were accepted. Events were defined as sleep spindle peak-locked 5 s long epochs (± 2.5 s centered on the spindle peak).

Ripples: The signal was first filtered in the range from 80-120 Hz and the analytical amplitude was extracted from a Hilbert transform in accordance with previously reported detection algorithms (Helfrich et al., 2019; Staresina et al., 2015). The analytical signal was smoothed with a 100ms window and z-scored. Candidate events were identified as epochs exceeding a z-score of 2 for at least 25ms and a maximum of 200ms and had to be spaced by at least 500ms. We determined the instantaneous ripple frequency by detecting all peaks within the identified segment. The identified events were time-locked to the ripple trough in a time window of ± 0.5 s. Overlapping epochs were merged. Epochs that contained IEDs or sharp transients were discarded.

Interictal epileptic discharge (IED) detection: We detected IEDs using automated algorithms on all channels located in the MTL. All cut-offs were chosen in accordance with recently published findings (Gelinas et al., 2016; Staresina et al., 2015) and were confirmed by a neurologist who visually verified the detected events. The continuous signal was filtered front and backwards between 25-80 Hz and the analytical amplitude was extracted from the Hilbert transform and then z-scored. Events were detected when this signal was 3 SD above the mean for more than 20ms and less than 100ms.

HFB, population activity and active periods analysis: The high frequency band (HFB) activity is typically defined from 70-180 Hz (Leszczyński et al., 2020). To avoid confounding true HFB activity with ripple-band activity (upper cut-off ∼120 Hz), we defined HFB activity as the average power in this frequency range from 120-180 Hz. The multi-taper spectral estimates where averaged into a single trace per electrode. The dynamics of the population activity were expressed as a population vector (Ebitz and Hayden, 2021). At every time point HFB activity was represented as a point *P* in a *n*-dimensional coordinate system where *n* reflects the number of electrodes. The population vector was then constructed by taking the Euclidean distance *d* between adjacent time points within in a given region-of-interest (ROI), hence, providing a single time course per ROI.

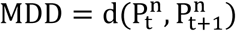

Active periods were defined as epochs where the smoothed (100ms window) HFB signal exceeded a z-score of 1 for at least 50ms (Grosmark et al., 2012).

#### Statistical analysis

Unless stated otherwise, we used cluster-based permutation tests (Maris and Oostenveld, 2007) to correct for multiple comparisons as implemented in FieldTrip (Monte Carlo method; 1000 iterations). Clusters were formed in time/frequency (e.g. **Fig. 1A/G-J**) or space (e.g. **Fig. 1D, 2C-E/G-H**) by thresholding dependent t-tests or linear correlations at p < 0.05. Correlation values were transformed into t-values using the following formula:

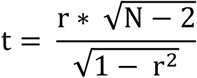

A permutation distribution was then created by randomly shuffling condition labels (paired t-tests) or subject labels (correlation). The permutation p-value was obtained by comparing the cluster statistic to the random permutation distribution. The clusters were considered significant at p < 0.05 (two-sided).

Effect sizes were quantified by means of Cohen’s d or the correlation coefficient rho. To obtain effect sizes for cluster tests, we calculated the effect size separately for all channel, frequency and/or time points and averaged across all data points in the cluster. Repeated-measures ANOVAs were Greenhouse-Geisser corrected. We further utilized linear mixed effect models with subjects as random intercepts.

**Fig. S1.**
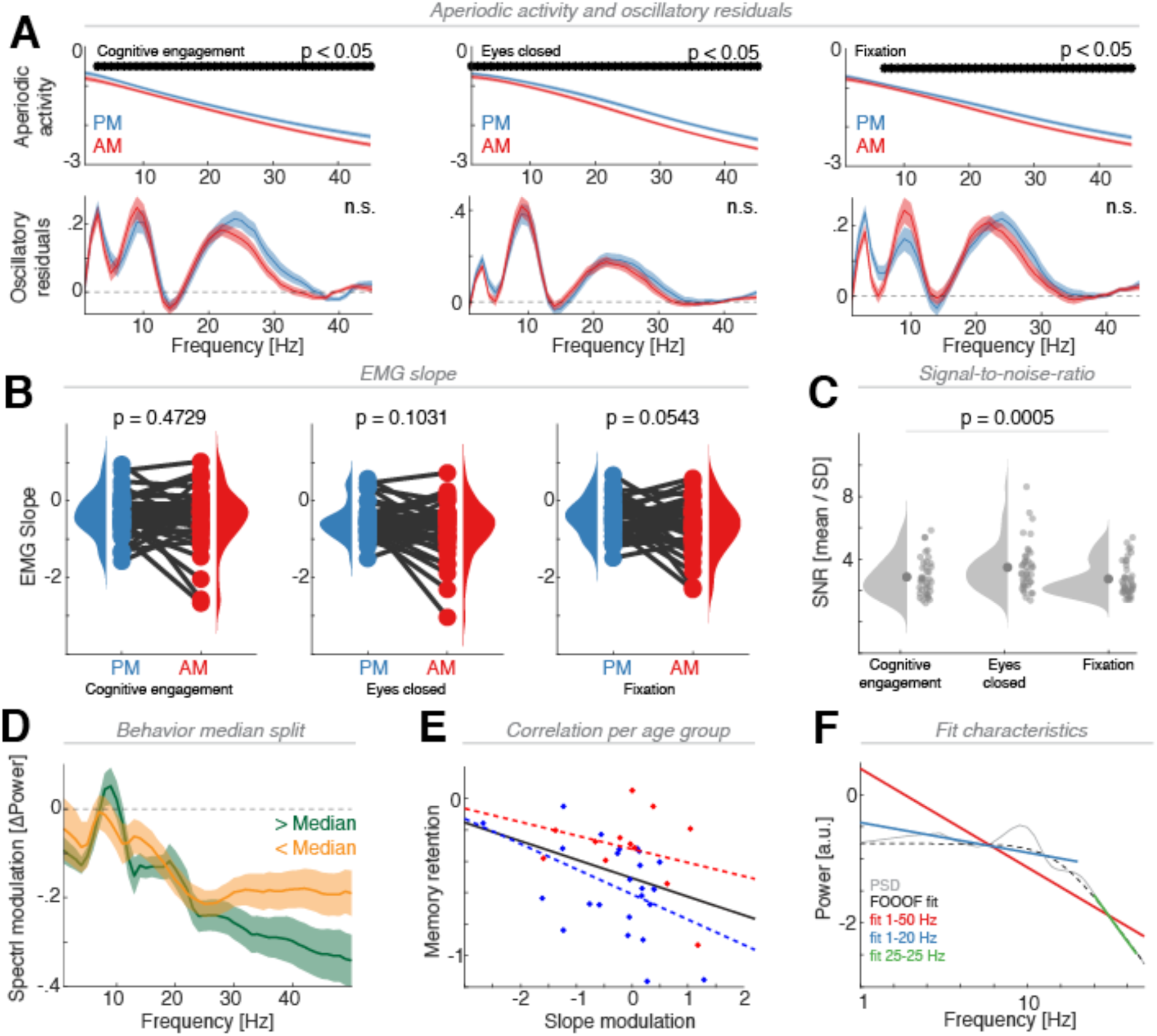
Control analyses related to Fig. 1. (**A**) Separation of aperiodic (upper row) and oscillatory activity (lower row) for all three resting state conditions (cognitive engagement, eyes closed, central fixation; visualized for electrode Cz). The modulation across a night of sleep was confined to aperiodic activity (fdr-corrected two-tailed t-tests) with consistently reduced power in the AM (red) as compared to the PM (blue) recordings in all three conditions. Note, this effect did not pertain oscillatory activity (lower row; all p-values > 0.05; n.s. = not significant). (**B**) Spectral slope of the EMG across all three resting state conditions shows no main effect of time-of-day. Results remained unchanged after partialling out the EMG slope from the correlational analyses. (**C**) Overall signal-to-noise ratio across all channels per participants reveals a main effect of condition, with the highest SNR observed for the eyes closed condition. (**D**) Median split of the behavioral memory retention metric to illustrate that subjects who performed better exhibited a larger broadband shift in the high frequency range. (**E**) Correlational analyses with the mean regression line (black) on the group level and separate fits for the young (red) and older (red) subjects. Note the general relationship of slope downscaling and behavior is preserved for older adults, but there is a shift along the y-axis, indicating that younger subjects performed better on average than older adults, as previously reported (Helfrich et al., 2018; Mander et al., 2013). (**F**) Illustration of suboptimal (red, linear fit 1-50 Hz; blue, linear fit 1-20 Hz) and optimal linear fits (green, 25-45 Hz), superimposed on an example spectrum (grey) that is characterized by both oscillatory bumps as well as 1/f background activity, which includes a knee (black dashed line indicates the appropriate FOOOF model).

**Fig. S2.**
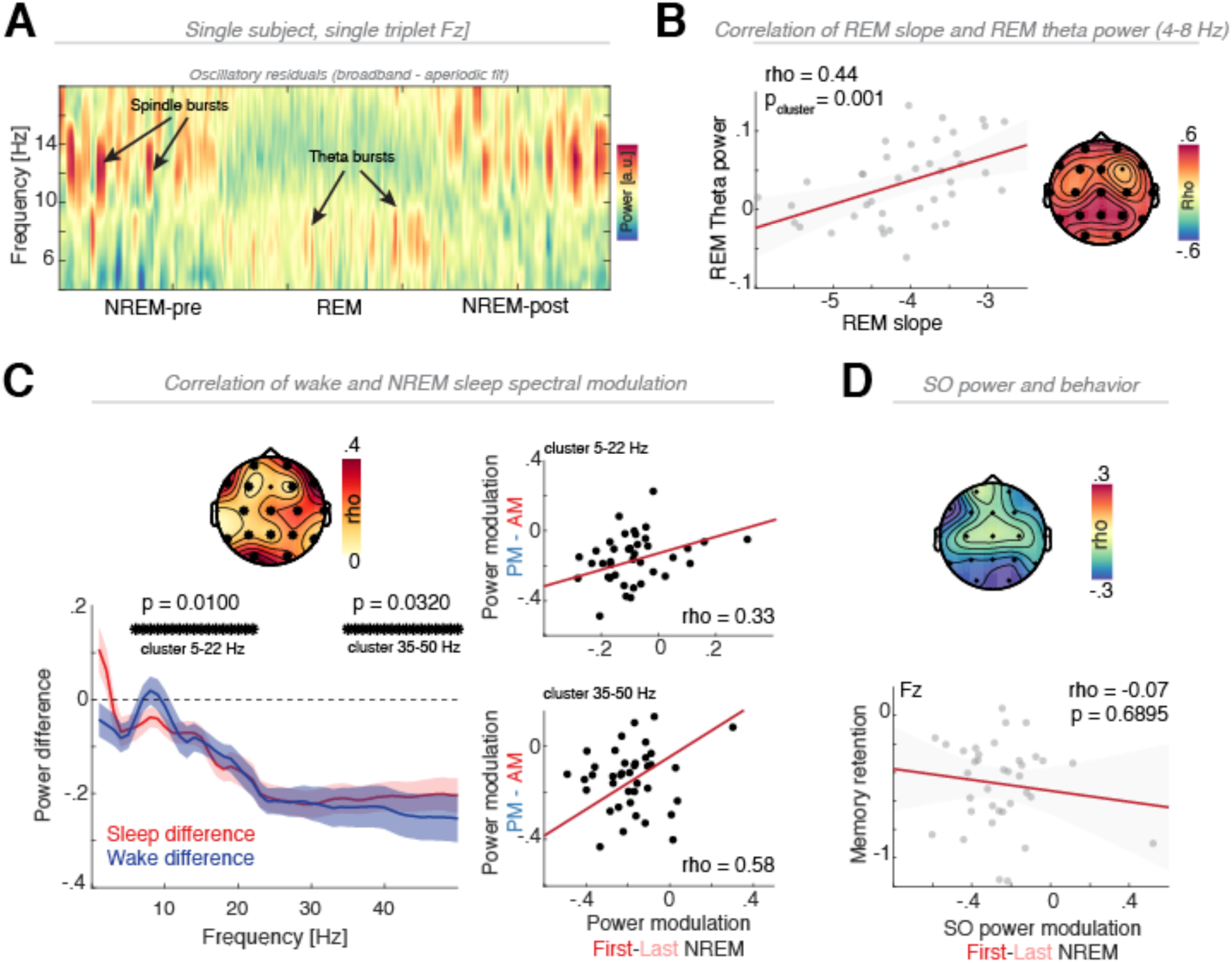
Control analyses related to Fig. 2. (**A**) Representative spectrogram of a single subject for a single NREM-REM-NREM triplet to highlight the presence of state-specific, short-lived spindle and theta burst. Note that the 1/f estimates have been discounted from the broadband spectrum to isolate the oscillatory residuals. (**B**) Correlation of REM slope and 1/f-corrected REM theta power reveals a widespread effect. Note that flatter REM slopes predict higher REM theta power. (**C**) Correlation of the spectral modulation in the wake state (PM-AM, cf. **Fig. 1A**) and NREM sleep (cf. **Fig. 2C**) indicates that the modulation as observed in one state is predictive of the other state (one outlier data point is not shown; correlations remained unchanged after accounting for this data point). (**D**) No significant correlation was observed for modulation of SO power across the night and behavior (cluster test; no significant clusters).

**Table S1.**

| Subject | Recording Duration | Total Sleep Time [h] | WASO [min] | N1 [min] | N2 [min] | SWS [min] | REM [min] |
| --- | --- | --- | --- | --- | --- | --- | --- |
| 1 | 8.0 | 6.1 | 89 | 11 | 200 | 63 | 92 |
| 2 | 8.0 | 6.0 | 111 | 31 | 209 | 51 | 72 |
| 3 | 7.9 | 6.0 | 102 | 36 | 194 | 79 | 52 |
| 4 | 7.8 | 2.1 | 321 | 33 | 30 | 37 | 24 |
| 5 | 7.9 | 6.6 | 69 | 21 | 270 | 1 | 108 |
| 6 | 7.4 | 5.6 | 82 | 19 | 222 | 40 | 58 |
| 7 | 7.9 | 5.3 | 139 | 44 | 198 | 12 | 65 |
| 8 | 7.9 | 3.9 | 195 | 29 | 111 | 68 | 24 |
| 9 | 7.9 | 6.9 | 50 | 22 | 145 | 110 | 138 |
| 10 | 8.0 | 7.2 | 40 | 16 | 225 | 111 | 81 |
| 11 | 8.1 | 6.2 | 101 | 25 | 189 | 105 | 52 |
| 12 | 7.7 | 4.4 | 187 | 27 | 169 | 40 | 27 |
| 13 | 7.8 | 5.4 | 132 | 16 | 164 | 82 | 61 |
| 14 | 7.9 | 7.0 | 41 | 15 | 236 | 57 | 112 |
| 15 | 8.0 | 5.8 | 121 | 24 | 200 | 70 | 53 |
| 16 | 7.9 | 6.9 | 47 | 24 | 309 | 2 | 81 |
| 17 | 8.0 | 6.5 | 71 | 21 | 224 | 57 | 87 |
| 18 | 7.9 | 4.9 | 38 | 11 | 147 | 71 | 66 |
| 19 | 7.7 | 4.9 | 157 | 21 | 198 | 15 | 62 |
| 20 | 7.9 | 4.2 | 162 | 11 | 114 | 108 | 21 |
| 21 | 8.0 | 6.4 | 57 | 20 | 169 | 123 | 73 |
| 22 | 7.9 | 6.2 | 88 | 15 | 260 | 59 | 40 |
| 23 | 8.0 | 6.0 | 100 | 17 | 196 | 100 | 46 |
| 24 | 8.0 | 3.9 | 230 | 22 | 144 | 43 | 27 |
| 25 | 8.0 | 6.9 | 59 | 31 | 230 | 39 | 114 |
| 26 | 7.9 | 6.7 | 57 | 18 | 232 | 33 | 118 |
| 27 | 8.0 | 7.1 | 44 | 30 | 187 | 137 | 71 |
| 28 | 8.0 | 7.7 | 6 | 6 | 230 | 123 | 105 |
| 29 | 8.0 | 6.2 | 93 | 10 | 200 | 118 | 43 |
| 30 | 8.0 | 7.2 | 31 | 14 | 198 | 141 | 77 |
| 31 | 8.0 | 7.1 | 30 | 14 | 203 | 154 | 57 |
| 32 | 8.0 | 7.6 | 10 | 18 | 213 | 85 | 142 |
| 33 | 8.0 | 7.2 | 16 | 10 | 286 | 61 | 76 |
| 34 | 8.0 | 7.4 | 18 | 6 | 159 | 162 | 120 |
| 35 | 8.0 | 7.2 | 31 | 28 | 184 | 107 | 114 |
| 36 | 8.0 | 6.8 | 42 | 11 | 184 | 144 | 68 |
| 37 | 8.0 | 6.9 | 21 | 19 | 181 | 128 | 85 |
| 38 | 8.0 | 7.6 | 10 | 16 | 209 | 106 | 126 |
| 39 | 8.0 | 7.7 | 4 | 6 | 215 | 123 | 120 |
| 40 | 8.0 | 7.1 | 19 | 17 | 223 | 90 | 98 |
| Group | 7.9 ± 0.1 | 6.2 ± 1.2 | 80.5 ± 68.5 | 19.6 ± 8.6 | 196.4 ± 49.0 | 81.4 ± 43.1 | 76.4 ± 33.2 |

#### Sleep metrics related to study 1

Participants went to bed at their habitual sleep time and were given the opportunity to sleep for up to 8h. Sleep staging was the carried out on the continuous epochs in 30s segments. Furthermore note that the rater flagged epochs of excessive noise as artifactual and hence, those epochs were excluded from staging.

**Table S2.**

| Subject | Recording Duration | Total Sleep Time [h] | WASO [min] | N1 [min] | N2 [min] | SWS [min] | REM [min] |
| --- | --- | --- | --- | --- | --- | --- | --- |
| 1 | 9.0 | 6.8 | 110 | 40 | 263 | 40 | 66 |
| 2 | 13.0 | 4.0 | 496 | 31 | 166 | 35 | 11 |
| 3 | 12.9 | 7.7 | 295 | 55 | 267 | 40 | 99 |
| 4 | 13.0 | 8.2 | 113 | 42 | 147 | 221 | 81 |
| 5 | 8.6 | 4.9 | 223 | 14 | 192 | 53 | 36 |
| 6 | 12.0 | 9.7 | 119 | 50 | 327 | 89 | 119 |
| 7 | 13.0 | 7.0 | 356 | 52 | 262 | 19 | 88 |
| 8 | 13.2 | 7.4 | 307 | 48 | 291 | 78 | 29 |
| 9 | 13.0 | 6.3 | 371 | 32 | 201 | 42 | 101 |
| 10 | 14.0 | 9.1 | 284 | 66 | 245 | 105 | 129 |
| 11 | 13.2 | 5.3 | 353 | 58 | 93 | 145 | 20 |
| 12 | 13.0 | 10.8 | 110 | 28 | 355 | 207 | 60 |
| 13 | 13.8 | 6.9 | 242 | 52 | 218 | 89 | 56 |
| 14 | 10.8 | 9.0 | 106 | 15 | 382 | 94 | 50 |
| 15 | 13.6 | 8.7 | 65 | 25 | 346 | 84 | 67 |
| Group | 12.4 ± 1.6 | 7.5 ± 1.9 | 236.7±128.6 | 33.6 ± 16.9 | 250.3 ± 82.4 | 89.4 ± 60.4 | 67.5 ± 35.4 |

#### Sleep metrics related to study 3

We aimed to include continuous ∼12-13h recording blocks, which were roughly obtained between 8pm and 8am. However, in some subjects the recording started later or stopped earlier due to ongoing testing or clinical considerations. Sleep staging was the carried out on the continuous epochs in 30s segments. Furthermore note that the rater flagged epochs of excessive noise as artifactual and hence, those epochs were excluded from staging. Note that wake-after-sleep-onset values (WASO) are inflated, given that the clinical routine often wakes patients around 6am and hence, the last two hours of the recording often have been spent awake.

